# IGT/LAZY family genes are differentially influenced by light signals and collectively required for light-induced changes to branch angle

**DOI:** 10.1101/2020.07.15.205625

**Authors:** Jessica Marie Waite, Christopher Dardick

**Affiliations:** United States Department of Agriculture (USDA) Appalachian Fruit Research Station, 2217 Wiltshire Road, Kearneysville, WV, U.S.A.; USDA Tree Fruit Research Laboratory, 1104 N Western Avenue, Wenatchee, WA, U.S.A

**Keywords:** IGT family, LAZY family, plant architecture, gravitropic set point angle, branch angle, light signaling

## Abstract

Plants adjust their growth orientations in response to environmental signals such as light and gravity in order to optimize photosynthesis and access to nutrients. However, given the fixed nature of gravity, understanding how light and gravity signals are integrated is challenging. Branch orientation, or gravitropic set point angle, is a key aspect of plant architecture, set with respect to gravity and shown to be altered by changes in light conditions. The IGT gene family, also known as the *LAZY* family, contains important components for branch angle and gravity responses, including three gene clades: *LAZY, DEEPER ROOTING (DRO)*, and *TILLER ANGLE CONTROL (TAC). LAZY* and *DRO* genes promote upward branch orientations downstream of amyloplast sedimentation, and upstream of auxin redistribution in response to gravity. In contrast, *TAC1* promotes downward branch angles in response to photosynthetic signals. Here, we investigated the influence of different light signaling pathways on *LAZY* and *DRO* gene expression, and their role in light regulation of branch angle responses. We found differential effects of continuous light and dark, circadian clock, photoreceptor-mediated signaling, and photosynthetic signals on *LAZY* and *DRO* gene expression. Phenotypic analysis revealed that *LAZY* and *DRO* genes are collectively required for branch angle responses to light.

**Highlight:** *LAZY* and *DRO* gene expression responds differentially to changes in light regime and signaling. Loss of multiple *LAZY* and *DRO* genes leads to loss of branch angle response to light.

## Introduction

Plant responses to light and gravity are crucial to their proper development and survival. When environmental changes occur such as light quality or the orientation of the plant with respect to gravity, the stimulus initiates a signaling cascade, relaying crucial information for the re-orientation of plant organs to ensure maximal access to light and soil resources. In this way, the overall shape, or architecture, of a plant is dependent on both light and gravity. A number of studies have shown that light and gravity pathways influence one another. For example, in the hanging plant *Tradescantia*, treatment with different inhibitors of photosynthetic activity led to a change in the gravitropic set point angle of the branches (Digby and Firn, 2002), and in maize, coleoptiles grown in a rotating clinostat to reduce perceived gravity show an enhanced phototropic response (Nick and Schäfer, 1988). In addition, recent studies performed in microgravity have begun to parse apart the individual influences of different light and gravity pathways on plant development and shape (Vandenbrink *et al*., 2014). Taken together, organ orientation is a consequence of the integration of light and gravity stimuli and has a profound influence on the development of plant shape.

One aspect of plant architecture that is strongly influenced by both light and gravity is lateral organ orientation, or the angle at which organs such as branches, lateral roots, and leaves grow. Optimizing lateral organ orientation can potentially benefit crops in terms of space and resource use, both above and below ground (Scorza *et al*., 2000; Uga *et al*., 2013; Li *et al*., 2017). Work in Arabidopsis has shown that light conditions can alter branch set point angles (Roychoudhry *et al*., 2017; Waite and Dardick, 2018), and leaf orientation is influenced by neighbor detection mechanisms using red/far-red light sensing in the leaf (Pantazopoulou *et al*., 2017). Taken together, organ orientation is a consequence of the integration of light and gravity stimuli and has a profound influence on the development of plant shape.

The IGT gene family, named for a highly conserved “IGT” amino acid motif, includes three distinct clades, *LAZY, TILLER ANGLE CONTROL (TAC)* and *DEEPER ROOTING (DRO)*, and plays a key role in the genetic control of lateral organ orientation (Li *et al*., 2007; Yu *et al*., 2007; Ku *et al*., 2011; Dardick *et al*., 2013; Dong *et al*., 2013; Uga *et al*., 2013; Yoshihara *et al*., 2013; Guseman *et al*., 2017). Recent studies have demonstrated that *LAZY* and *DRO* genes influence organ orientation downstream of amyloplast sedimentation and upstream of auxin localization (Taniguchi *et al*., 2017; Yoshihara and Spalding, 2017; Nakamura *et al*., 2019). In the roots, the “anti-gravitropic” phenotype of a *lazy dro* quadruple mutant requires functional statoliths, and in the shoot requires functional endodermal cells (Kawamoto *et al*., 2020). Mechanistic studies on DRO1 have shown that the protein is localized to the nucleus under native conditions in roots tips. However, upon reorientation of roots, the protein becomes polarly localized along the lower plasma membrane of the gravity sensing columella cells prior to subsequent changes in PIN localization (Waite *et al*., 2020; Furutani *et al*., 2020).

Recently, we showed that the expression of another IGT gene family member, *TAC1*, is influenced by photosynthetic signals and contributes to branch orientation (Waite and Dardick, 2018). Here, we seek to understand whether different aspects of light signaling also affect *LAZY* and *DRO* gene expression. We demonstrate through mutant analysis, light, and chemical treatments that photoreceptor-mediated light signaling pathways, photosynthetic signals and photosynthate, and the circadian clock have differential influences on *LAZY* and *DRO* gene expression. We find that changes in gravitropic setpoint angles in response to light requires the presence of multiple IGT genes. Our work suggests the existence of both IGT-dependent and independent pathways for branch angle response to changing light regimes. We conclude that IGT genes are differentially regulated by light signals, and that *LAZY* and *DRO* genes are required for light-induced changes to branch growth angles. We hypothesize that members of the IGT gene family serve as integrators of environmental signals regulating plant architecture.

## Materials and Methods

### Plant Materials

The Columbia (Col-0) ecotype was used as wild-type in all experiments. The *lazy1* mutant used here was a T-DNA insertion line (GABI_591A12) obtained from NASC (https://arabidopsis.info). The *tac1 lazy1* double mutant was generated by crossing the *tac1* and *lazy1* single SALK mutants (*tac1*, T-DNA insertion line CS825872 from ABRC) as described in Hollender et al 2020. *lazy6* and *lazy1 lazy6* lines were generated by transforming *LAZY6* CRISPR constructs into Col-0 and *lazy1* backgrounds, respectively. Briefly, for CRISPR constructs, the *LAZY6* target sequence was identified using crispr-plant (http://www.genome.arizona.edu/crispr/), and cloned into the pHEE401E vector (Wang *et al*., 2015) using Gibson cloning (Gibson *et al*., 2009). The resulting *LAZY6* CRISPR line shown here contains a single G insertion at nucleotide position 229 of the cDNA, resulting in a premature stop after the 80^th^ codon, and was phenotypically representative of multiple CRISPR mutants. The *35S::DRO1* and *35S::LAZY1* constructs were cloned as preciously described (Guseman *et al*., 2017; Hollender *et al*., 2020). Similarly to these, the *35S::LAZY6* construct was cloned by ligating the *LAZY6* (At3g27025) CDS into the multiple cloning site downstream of the 35S promoter of a modified pBIN-AFRS expression vector (Belknap *et al*., 2008). All three constructs were transformed into *lazy1* mutant Arabidopsis plants via floral dip (Clough and Bent, 1998). Signaling mutants *phyA phyB* and *phyA phyB phyD phyE* (Hu *et al*., 2013), *cry1 cry2* (Mockler *et al*., 1999), *phot1 phot2* (Kinoshita *et al*., 2001), *cop1-6* (Ang and Deng, 1994), *pif1 pif3 pif4 pif5* (Lilley *et al*., 2012), and *hy5 hfr1 laf1* (Jang *et al*., 2013) were previously described. For seedling expression studies, seeds were surface sterilized and sown on square plates containing half strength Murashige and Skoog (MS) medium and 0.8% bactoagar, and grown vertically, following our standard lab practice. Once sown, seedlings were stratified at 4 °C in the dark for 2 d, then placed in growth chambers at 20 °C with a 16 h light/8 h dark photoperiod (∼100 µmol m–2 s–1). For adult phenotyping studies, 14-day old seedlings were transplanted to soil and allowed to grow in these conditions until bolting.

### RNA extraction and qPCR

Seedlings were grown on vertical plates for 10–14 d. Four biological replicates were used. Each biological replicate consisted of a plate of 10–12 seedlings. RNA was extracted using a Directzol RNA Extraction Kit (Zymo Research, http://www.zymoresearch.com). qPCR was performed as previously described by Waite and Dardick (2018). Briefly, each reaction was run in triplicate using 50 ng of RNA in a 12 µl reaction volume, using the Superscript III Platinum SYBR Green qRT-PCR Kit (ThermoFisher Scientific, https://www.thermofisher.com). The reactions were performed using an ABI7900 qPCR machine (Applied Biosystems, now ThermoFisher Scientific, https://www.thermofisher.com). Quantification of Arabidopsis samples was performed using a standard curve derived from a serially diluted standard RNA run in parallel. UBC21 was used as an internal control to normalize expression in light experiments as it was identified as a highly constitutive gene (Czechowski *et al*., 2005), and IPP2 was used for circadian experiments, as described previously by Imaizumi et al. (Imaizumi *et al*., 2005).

### Branch angle phenotypes

For shoot branch phenotypes, seedlings were grown for 2 weeks on plates, then transplanted into 4 inch pots containing Metromix 360 soil (Sun-Gro Horticulture, http://www.sungro.com) and grown until bolting (∼15–18 cm in height). Plants were then transferred to continuous light or dark conditions for 72 h. Images were taken using a Canon EOS Rebel T3 camera (http://global.canon/en/index.html, last accessed 27 July 2018).

### Light and time-course experiments

For seedling light experiments, plants were grown for 10 days on vertical plates in 16:8 h long day light conditions in a growth chamber before transfer to experimental light conditions. For comparisons between light and dark, plates were moved to chambers with either continuous light or continuous dark conditions for 72 h, then whole seedlings were collected and flash frozen at 10.00am [Zeitgeber time (ZT4)]. For comparisons between light colors, plates were moved to chambers with continuous white (W), R (660 nm), blue (B; 480 nm), or FR light (738 nm) for 72 h and whole seedlings were collected at 10.00am (ZT4). Matching growth chambers fitted with white, red, blue, and far-red LED lamps from PARsource (http://parsource.com) were used for light color experiments. For circadian experiments, seedlings were grown for 10 d in 12L:12D light cycles, then transferred to continuous light and collected every 4 h for 84 h. For adult phenotypes and expression studies, plants were grown on soil for 5–6 weeks, until bolts reached 15–18 cm in height. Then plants were transferred to continuous light or dark conditions for 72 h, then imaged. For identification of light-related cis-elements in the IGT gene promoters, we used the AGRIS AtCis Database (https://agris-knowledgebase.org/AtcisDB/).

### Chemical treatments

For sucrose experiments, plants were germinated and grown on half-strength MS plates for 10 d, then transplanted to plates containing 1% sucrose. Plates were then moved to continuous light or dark conditions for 72 h and collected at 10.00am (ZT4). For photosynthesis experiments, plants were grown on vertical MS plates for 7 d, then transplanted onto media containing either Norflurazon (NF) (5 µM), DCMU (10 µM), Paraquat (PQ) (1 µM), or water (mock). Plates were then moved to continuous light or dark conditions for 72h and collected at 10.00 h (ZT4).

## Results

### Dark and light conditions influence on *LAZY* and *DRO* gene expression dynamics

The reported connections between light and organ orientation (Digby and Firn, 2002; Roychoudhry *et al*., 2017), and our recent finding that *TAC1* is a target of photosynthetic signals regulating branch angle (Waite and Dardick, 2018), led us to examine the relationship between light and the other IGT-family genes within the *LAZY* and *DRO* clades (Hollender and Dardick, 2015; Guseman *et al*., 2017). We first used a cis-element database (AGRIS AtCisDB, https://agris-knowledgebase.org/AtcisDB/) to identify light-, circadian- and auxin-related cis-elements in the promoters of *LAZY* and *DRO* genes (Supplementary Fig. S1). All IGT genes contained light related elements. GATA-box-containing cis-elements, which have been shown to be associated with light and circadian responsiveness (Teakle *et al*., 2002; Hudson and Quail, 2003), appeared in all examined promoters. *LAZY1, DRO2, DRO3*, and *LAZY6* contained T-box elements, which were identified to positively modulate light activation of a nuclear gene encoding a chloroplast protein (Chan *et al*., 2001). *DRO1* and *DRO2* promoters contained an Auxin Response Element, known for ARF binding (Ulmasov *et al*., 1999), and *DRO2* and *LAZY6* contained AtMyc2 sites, which have been implicated in blue light signaling and interaction with GATA-containing promoters (Yadav *et al*., 2005; Gangappa and Chattopadhyay, 2013). Cis-elements associated with phyA signaling (SORLREP and SORLIP) appeared in *DRO1, DRO3*, and *LAZY6*, and *LAZY6* additionally contained two G-box elements and a TGA1 site, both of which bind bZip transcriptions factors involved in light signaling (Schindler *et al*., 1992; Hudson and Quail, 2003).

We next designed a set of experiments to measure changes in IGT gene expression in response to different light regimes and signaling pathways. We were able to design reliable and efficient quantitative PCR primers for four of the *LAZY* and *DRO* clade genes: *LAZY1* (At5g14090), *LAZY6* (At3g27025), *DRO1* (At1g72490), and *DRO2* (At1g19115) (Supplementary Table S1) (Ge and Chen, 2016; Taniguchi *et al*., 2017; Yoshihara and Spalding, 2017; Guseman *et al*., 2017). Unfortunately, after testing multiple primer pairs, we were unable to find efficient quantitative PCR primers that could consistently detect *DRO3* (At1g17400) (data not shown). To minimize environmental variables, we measured expression responses for most light treatments using populations of 2 week-old seedlings grown on agar plates, in which *LAZY* and *DRO* gene expression has been determined (Yoshihara *et al*., 2013; Yoshihara and Spalding, 2017; Guseman *et al*., 2017; Hollender *et al*., 2020).

To understand both endpoint and dynamic responses of *LAZY* and *DRO* gene expression to continuous dark and light treatments, we performed 3-day time course experiments with sample collection times chosen to distinguish between fast (minutes to hours) and slow (days) responses (Fig. 1). Seedlings grown in 16:8 long-day conditions were transferred to continuous darkness for 3 days, and sampled at 0, 0.5, 1, 2, 4, 8, 12, 24, 48, and 72 hours after the change (Fig. 1A). A second set of seedlings, were grown in long-day conditions, transferred to continuous dark for 3 days, then returned to continuous light and sampled at the same intervals (Fig. 1B). We found that expression of the *LAZY1* gene was relative stable throughout both treatments, showing expression changes less than 2-fold among all samples. In contrast, *LAZY6* expression decreased significantly between 2-4 hours in the dark and remained low while grown in the dark. In seedlings returned to the light, expression peaked at 2 hours, then decreased and remained stable for the rest of the experiment. *DRO1* also decreased in the dark, although to a lesser degree. Upon transfer back to light conditions, expression was variable, with a peak at 2 hours. *DRO2* exhibited an increase of expression after 72 hours in darkness. Subsequent light treatment resulted in a sharp decrease over the first two hours, and expression remained low for the duration of the experiment. These data demonstrate that *LAZY* and *DRO* genes are light responsive and have differential dynamics in response to continuous dark and light treatments. The results led us to explore different aspects of light signaling – circadian rhythms, photoreceptor-mediated signaling, and photosynthesis – to determine the influence of each on *LAZY* and *DRO* gene expression.

**Figure 1:**
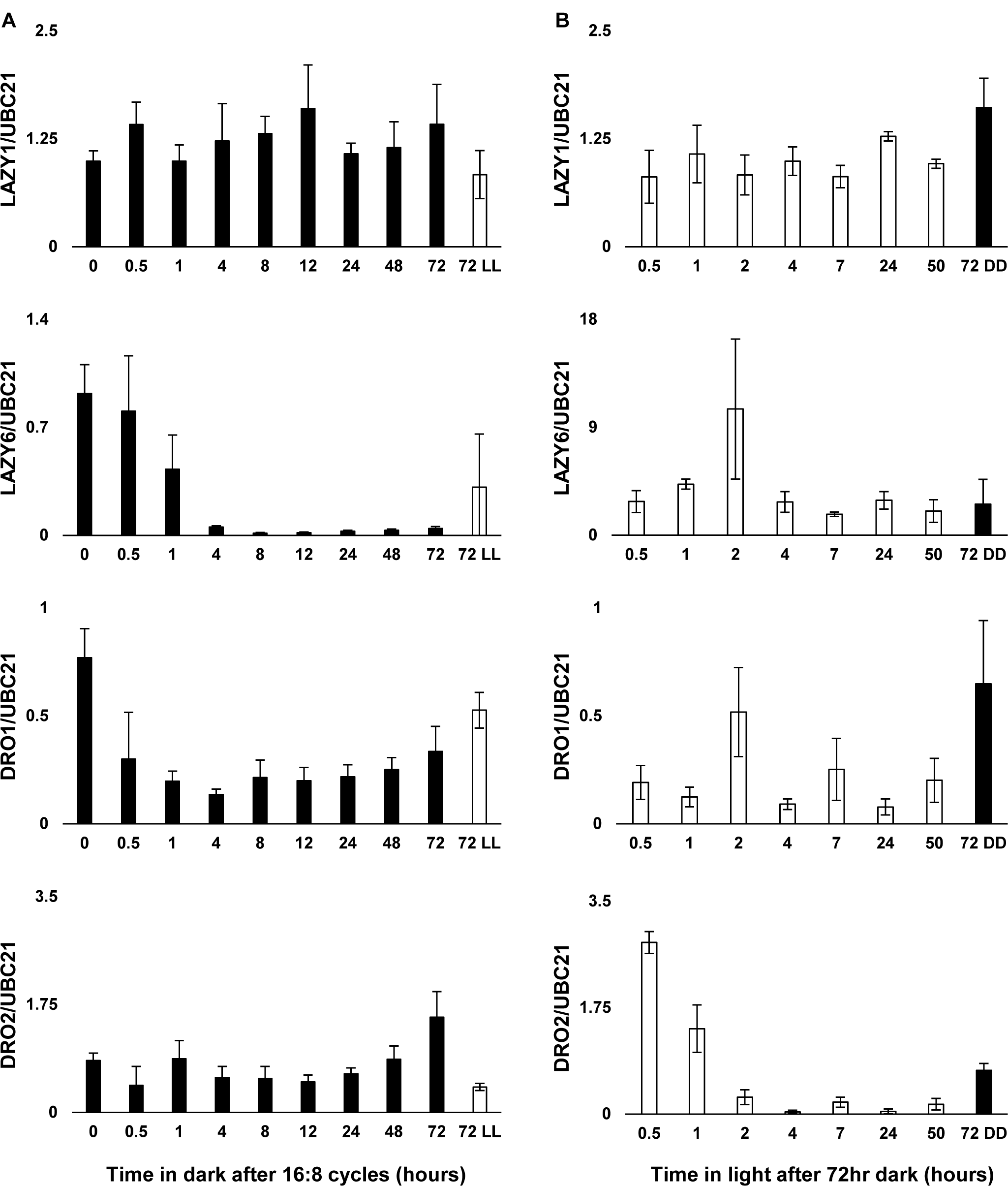
*LAZY* and *DRO* genes exhibit differential gene expression dynamics in response to continuous light and dark. A. Time-course of gene expression for *LAZY1, LAZY6, DRO1*, and *DRO2* from seedlings grown in 16:8 light cycles and then moved to continuous dark for 72 hours, relative to *UBC21* (Czechowski *et al*., 2005). For comparison, seedlings kept in 72 hours of continuous light were also included (far right). B. Gene expression time-course from seedlings grown in 16:8 light cycles, moved to continuous dark, then back to continuous light for 72 hours. For comparison, seedlings maintained in continuous darkness were also included (far right). Error bars represent SD between 4 biological replicates of 10-12 pooled seedlings per time point.

### Circadian influence on *LAZY* and *DRO* gene expression dynamics

The gene expression profiles we observed under continuous light or dark, particularly for *LAZY6* and *DRO1*, hinted at a potential role for circadian regulation in IGT gene expression. To test this, seedlings were entrained to 12:12 light/dark cycles, then transferred to continuous light conditions and sampled every 4 hours for 3.5 days (Fig. 2). *LAZY1, LAZY6*, and *DRO1* were consistent with circadian regulation, with peaks in expression occurring every ∼24 hours and shifting slightly over time (Fig. 2). In contrast, *DRO2* showed a weaker pattern, and instead appeared to steadily drop throughout the duration of the continuous light treatment.

**Figure 2:**
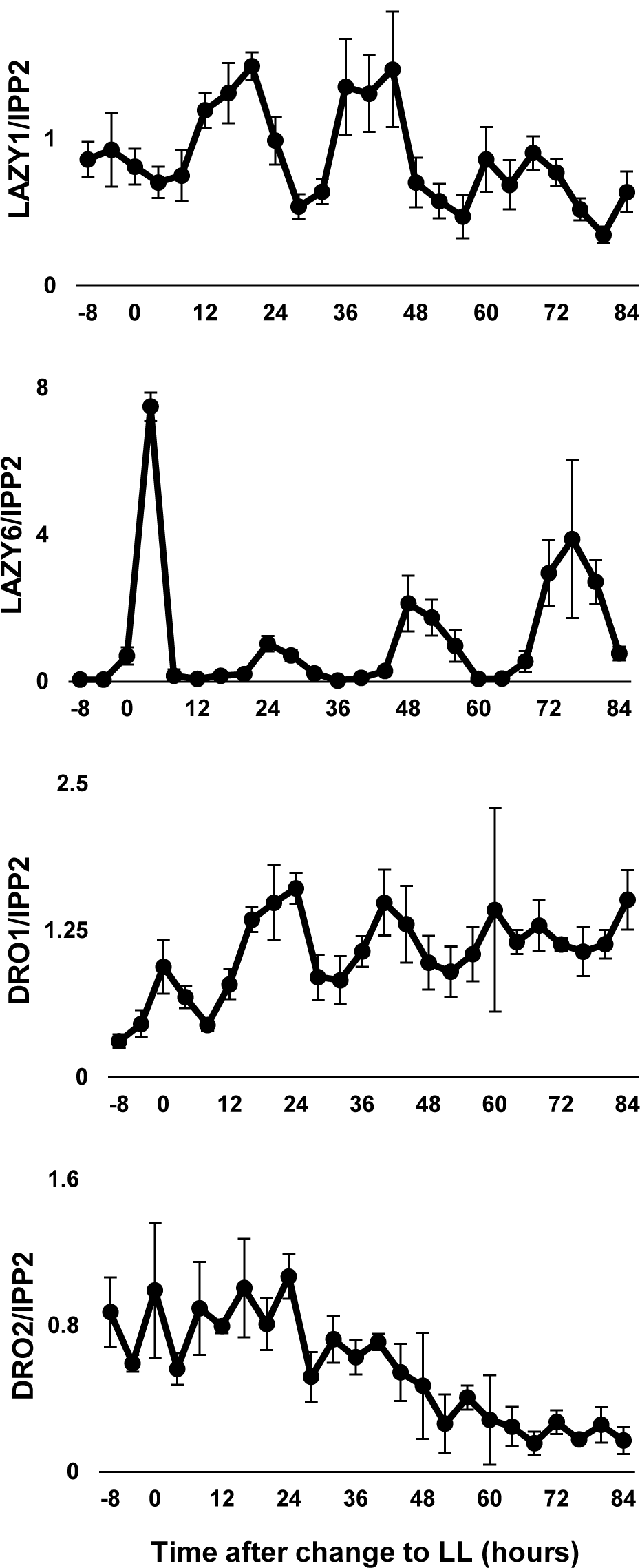
Gene expression of some IGT genes shows circadian rhythms. Time course of *LAZY1, LAZY6, DRO1*, and *DRO2* gene expression from seedlings entrained to a 12:12 light cycle, then moved to continuous light for 84 hours, relative to *IPP2* expression (Imaizumi *et al*., 2005). *LAZY1, LAZY6*, and *DRO1* show signatures of circadian rhythms, while *DRO2* steadily tapers with longer light exposure. Error bars represent SD between 4 biological replicates of 10-12 pooled seedlings per time point.

### *LAZY* and *DRO* gene expression responds differentially to light color and photoreceptors

To assess the effect of different light colors and photoreceptor-mediated pathways on the expression of *LAZY* and *DRO* genes, we began by growing seedlings in continuous Blue (B), Red (R), or Far-Red (FR) light for three days (Fig. 3A). A change from 16:8 White (W) light to B light had no significant effect on *LAZY1, DRO1*, or *DRO2*. In contrast, *LAZY6* expression was upregulated more than 3-fold. R light had a significant effect on all genes, leading to an increase in *LAZY1* expression, but a decrease in the other three genes. Finally, FR light only had a significant effect on the *DRO* genes, leading to a significant decrease in *DRO1* and *DRO2* expression (Fig. 3A). Together, this demonstrated a differential response of *LAZY* and *DRO* gene expression to different light colors.

**Figure 3:**
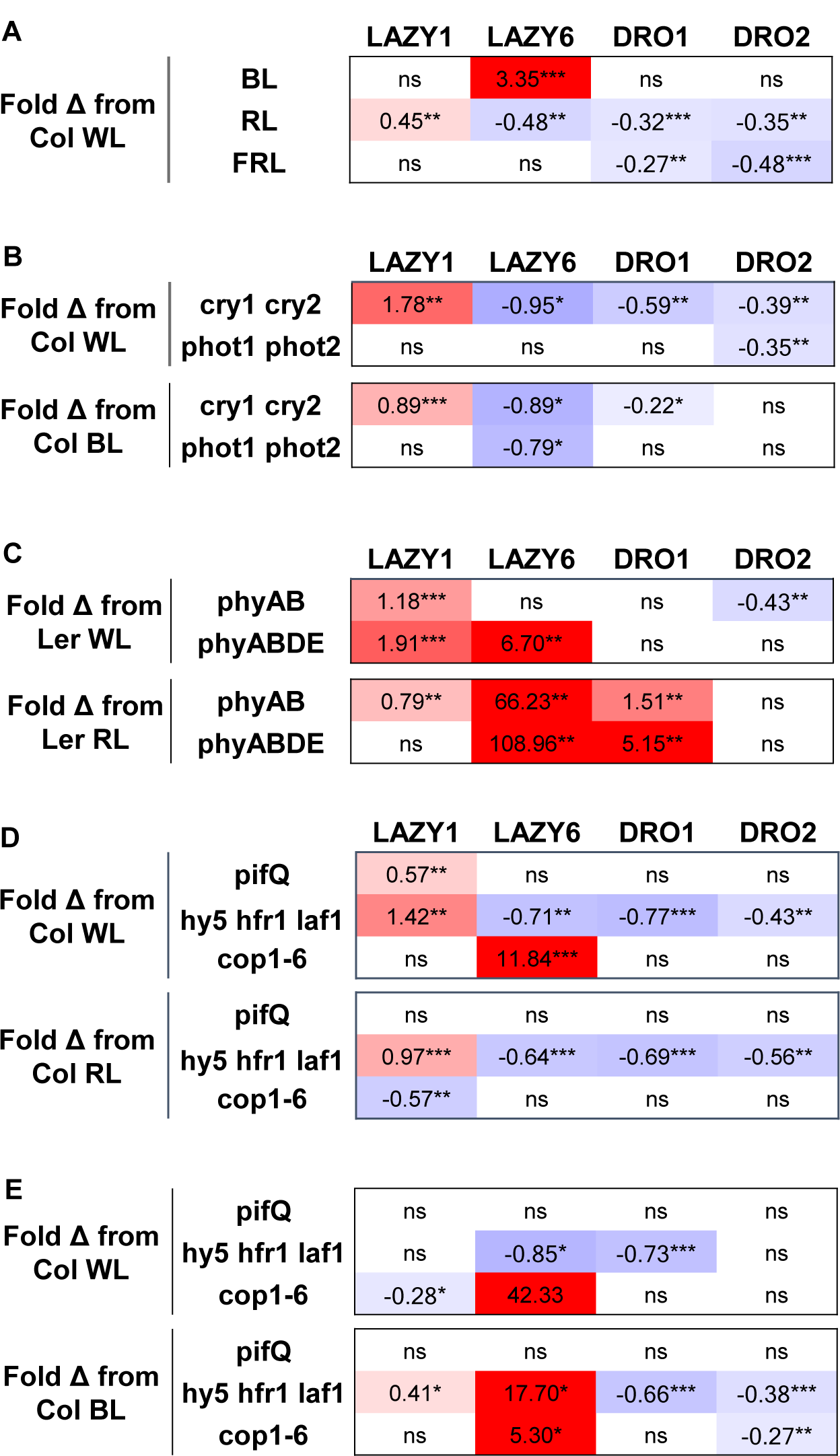
Effect of light color and photoreceptor-mediated signaling mutants on *LAZY* and *DRO* gene expression. A. Difference in expression between seedlings moved from 16:8 light cycles into continuous WL, and those moved into continuous B, R, or FR light. Significant fold changes from WL treatment are reported. B. Expression differences between Columbia wild type and blue light signaling mutants, *cry1 cry2* and *phot1 phot2*, moved from 16:8 light cycles to either continuous WL or BL. Significant changes from Col in WL or in BL are reported. C. Expression differences between Columbia wild type and red light signaling mutants, *phyAB* or *phyABDE*, moved from 16:8 light cycles to either continuous WL or RL. Significant changes from Col in WL or in RL are reported. D. Expression differences between Columbia wild type and light signal integration mutants, *pifQ, hy5 hfr1 laf1*, and *cop1-6*, moved from 16:8 light cycles to either continuous WL or RL. Significant changes from Col in WL or in RL are reported. E. Expression differences between Columbia wild type and light signal integration mutants, *pifQ, hy5 hfr1 laf1*, and *cop1-6*, moved from 16:8 light cycles to either continuous WL or BL. Significant changes from Col in WL or in BL are reported. Expression values were calculated relative to *UBC21*, and represent the average between 4 biological replicates of 10-12 pooled seedlings per comparison. Significant changes are color coded for direction and amplitude of expression change. * P<0.05, ** P<0.01, *** P<0.001.

To parse the regulatory influence of different photoreceptor-mediated pathways, we looked at expression in signaling mutant backgrounds associated with cryptochrome (cry), phototropin (phot), and phytochrome (phy) pathways, as well as downstream signaling genes known to integrate different light signals. Mutants were grown in continuous W, B, or R light for 3 days. While *LAZY1* showed no major change in expression in response to blue light alone, its expression significantly increased in a *cry1 cry2* background, both in white and blue light treatments, suggesting a repressive effect of the cryptochromes (Fig. 3B). The loss of *phot1* and *phot2*, however, appeared to have no effect on expression. *DRO1* and *DRO2* expression decreased in a *cry1 cry2* background (Fig. 3B). Similar to *LAZY1, DRO1* showed no significant differences between wild-type and *phot* double mutants, while *DRO2* expression decreased in the *phot* mutant background. Consistent with a greater B light-induced increase in *LAZY6* expression, downregulation was observed in both *cry* and *phot* mutant background under most treatments (Fig. 3B). This may suggest a positive role of these genes in regulating *LAZY6* expression.

Loss of two (*phyA* and *phyB*) or four *phys* (*phyA, B, D*, and *E*) had a largely positive effect on *LAZY* and *DRO* genes (Fig. 3C). *LAZY6* showed the most dramatic increases, between 6.7 and 108.9 fold in response to the loss of the *phy* genes. *LAZY1* increased significantly in the W light treatment, but to a lesser extent under R light conditions. *DRO1*, in contrast, showed no response to the absence of functional *phy* genes in W light conditions, but dramatic increases under R light. *DRO2* exhibited minor changes in expression, only significantly downregulated in the *phyA phyB* background in W light.

A number of genes are known to act downstream of multiple light signaling pathways, acting as a signal integration center (Paik *et al*., 2017; Podolec and Ulm, 2018; Xu, 2019). These include *PHYTOCHROME INTERACTING FACTORS* (*PIFs*), *ELONGATED HYPOCOTYL 5* (*HY5*), (*HFR1*), *LONG AFTER FAR-RED LIGHT 1* (*LAF1*), and *CONSTITUTIVE MORPHOGENESIS 1* (*COP1*). Similar to the photoreceptors, mutations in these genes showed differential effects on *LAZY* and *DRO* gene expression (Fig. 3D and 3E). *LAZY1* increased significantly under W light conditions in the *pifQ* mutant, which combines loss of four of the *PIF* genes (*PIF1,3,4,5*). However, this quadruple mutant had no significant effect on the expression of the other genes. *LAZY1* also showed upregulation in the triple *hy5 hfr1 laf1* mutant background under most conditions, and downregulation in the *cop1-6* mutant under some W and R light conditions. *LAZY6* showed the most dramatic changes overall. In the *hy5 hfr1 laf1* mutant background, *LAZY6* showed downregulation in W and R light, but a dramatic upregulation under B light. In a *cop1-6* background, *LAZY6* expression showed upregulation under W and B light, but this was lost under R light. *DRO1* was downregulated in the *hy5 hfr1 laf1* mutant under all conditions tested, and to a lesser extent, *DRO2* was also decreased in most light treatments. Together, these data suggest a strong influence of photoreceptors on *LAZY6*, a largely positive effect of *phy* loss and negative effect of *cry* loss on the *LAZY* and *DRO* genes. The triple *hy5 hfr1 laf1* mutant had a differential influence on gene expression changes for all genes in nearly all conditions, and *cop1* had large positive effects on *LAZY6*.

### Photosynthetic signals influence *LAZY* and *DRO* gene expression differentially

Light information from photosynthesis can also play an important regulatory role influencing branch angle, as we recently found with *TAC1* (Waite and Dardick, 2018). To initially assess how *LAZY* and *DRO* genes respond to inhibition of photosynthesis, we treated plants with 3 different chemical inhibitors (Fig. 4). Norflurazon (NF) inhibits carotenoid biosynthesis, 3-(3,4-dichlorophenyl)-1,1-dimethylurea (DCMU) blocks Photosystem II (PSII) function, and paraquat (PQ) inhibits PSI. Similar to light treatments, we transferred 2 week-old seedlings onto plates containing these chemicals and allowed them to grow in continuous light or dark for 3 days. Growth in the presence of NF had no effect on *LAZY1, LAZY6*, or *DRO1*, however a small but significant decrease could be seen for *DRO2* expression in dark-grown plants (Fig. 4A). Light grown plants in the presence of DCMU showed a decrease in expression of both *LAZY* genes and increased expression of both *DRO* genes (Fig. 4C). A decrease in *LAZY6* was also observed in dark grown plants. Treatment with PQ followed no identifiable pattern, with a decrease in *LAZY1* in dark grown plants, and an increase in *LAZY6* and decrease in DRO2 in the light (Fig. 4B). Treatment with photosynthate, in the form of exogenous sucrose, had very little effect on most genes, with the exception of *LAZY6*. The loss of *LAZY6* in darkness was rescued by growth in the presence of sucrose (Fig. 4D).

**Figure 4:**
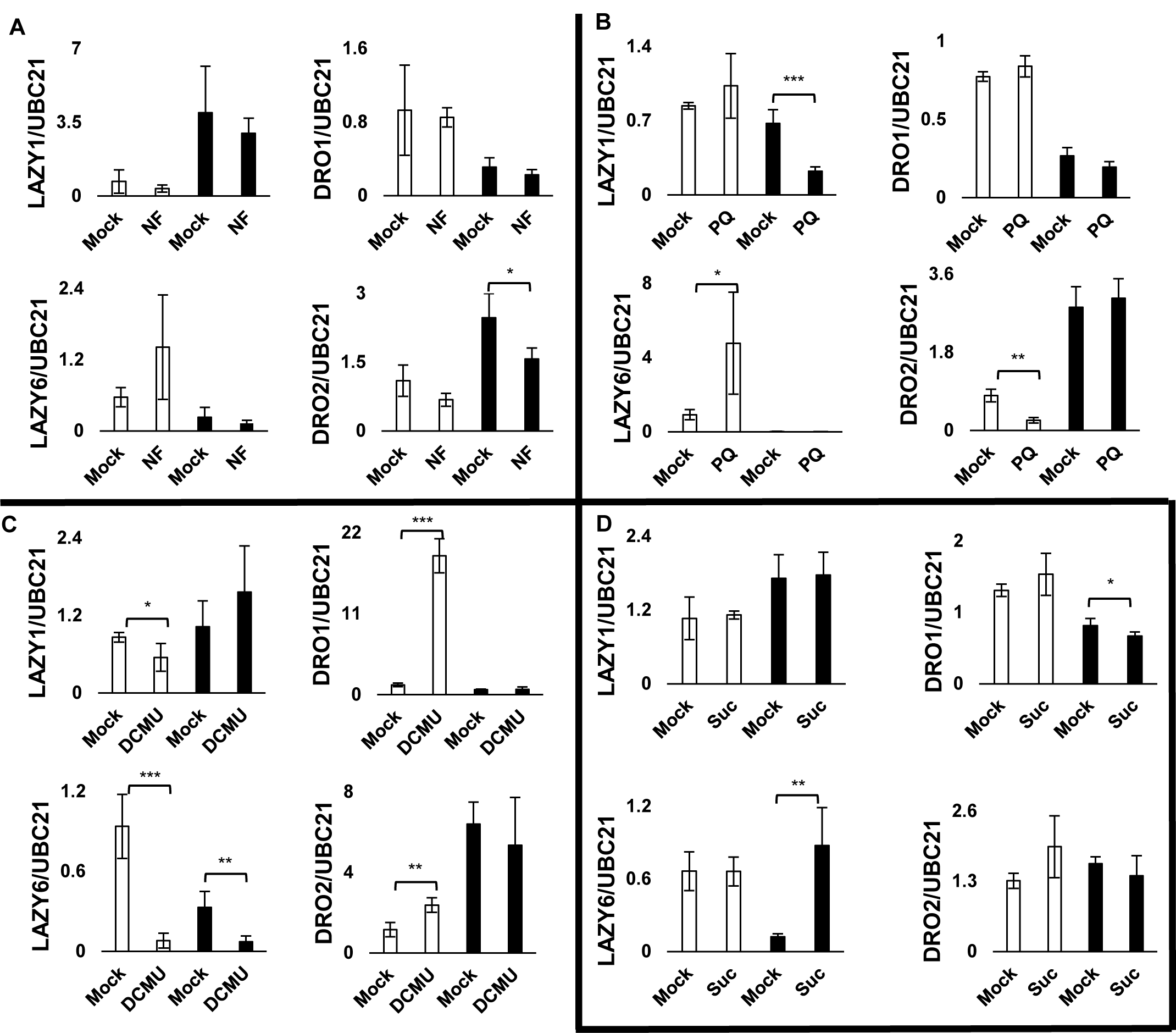
Photosynthesis inhibitors and sucrose have differential effects on IGT genes. Relative expression of *LAZY1, LAZY6, DRO1*, and *DRO2* in seedlings transplanted onto media containing either a mock treatment or A. Norflurazon, B. Paraquat, C. DCMU, or D. Sucrose and grown in continuous light or dark for 72 hours. Significant differences from mock treatments are reported. Expression values were calculated relative to *UBC21*. Error bars represent SD between 4 biological replicates of 10-12 pooled seedlings per time point. * P<0.05, ** P<0.01, *** P<0.001.

### Phenotypic responses of *LAZY* and *DRO* mutants to changes in light

Together, our gene expression data revealed a complex picture regarding the role of light in the regulation of IGT-family genes, with different light conditions and pathways affecting each *LAZY* or *DRO* gene differentially. To address the potential functional implications of light regulation on IGT genes in plant branch angle phenotypes, we designed a set of experiments to look at both mutant and overexpression phenotypes grown under different light environments. First, we grew Arabidopsis wild-type, *lazy1* single, *tac1 lazy1* double, and *lazy1 dro1 dro3* triple mutants under continuous light and continuous dark regimes (Taniguchi *et al*., 2017; Yoshihara and Spalding, 2017; Hollender *et al*., 2020). Adult plants were grown in long-day conditions until they began initiating branches, and then were moved into continuous light or continuous dark. As seen previously, we found that under continuous dark conditions wild-type branch tip angles narrowed and branches began to grow upward, against the gravity vector (Fig. 5A, and Waite and Dardick, 2018). In contrast, branch tips continued to grow at a wider, more horizontal angle under continuous light (Fig. 5A). We initially assessed these phenotypes up to 72 hours, the period after which we previously assessed gene expression, but found little difference in light-response phenotypes between 48 and 72 hours (Supplementary Fig. S2).

**Figure 5:**
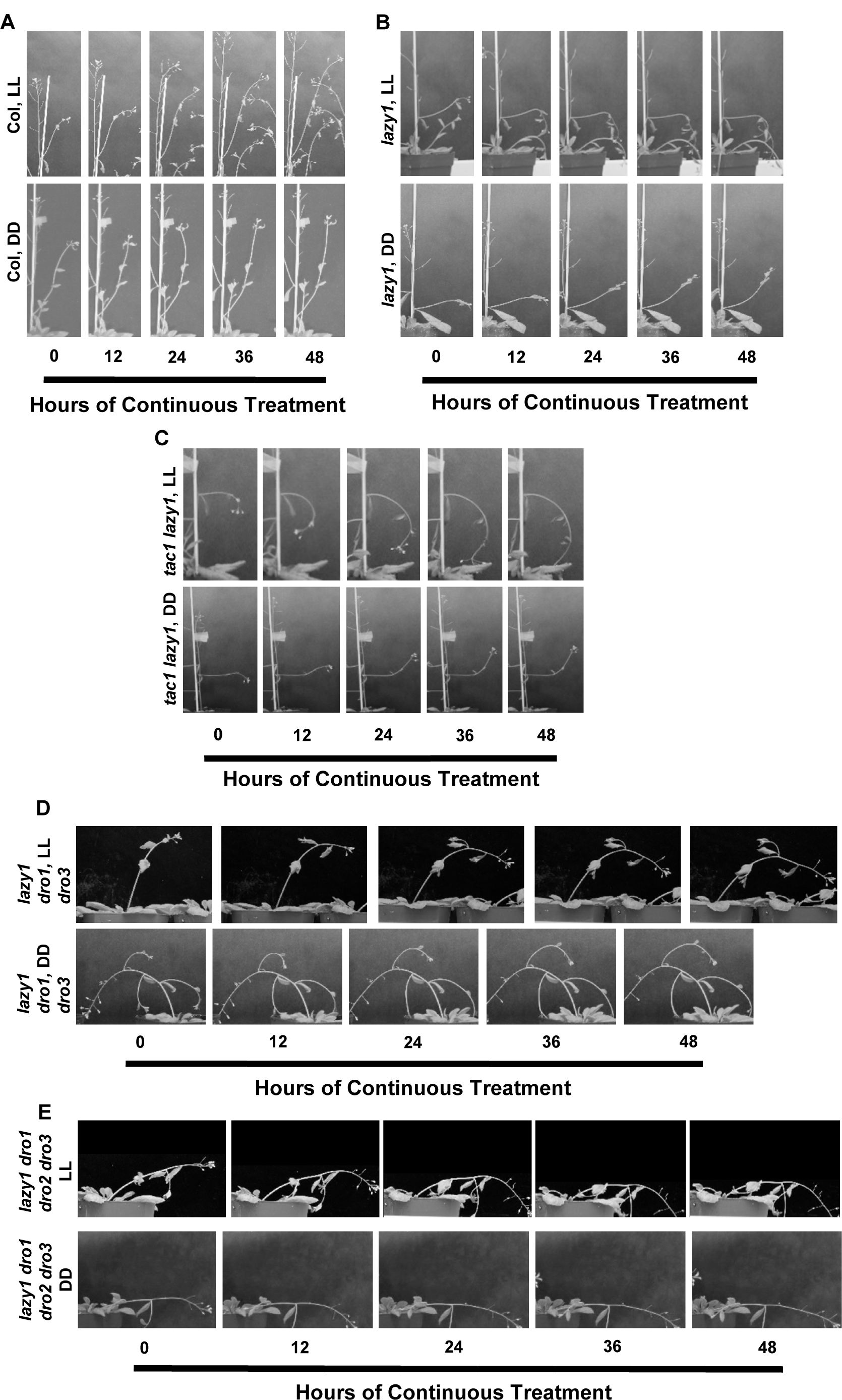
Loss of multiple *LAZY* and *DRO* genes confers phenotypic insensitivity to changes in light. Time course of A. Columbia wild-type, B. *lazy1* mutant, C. *tac1 lazy1*, D. *lazy1 dro1 dro3* triple mutant, and E. *lazy1 dro1 dro2 dro3* quadruple mutant branch angle response to continuous light and dark. Adult plants were grown in 16:8 light cycles and moved to continuous light or dark. Panels demonstrate branch and tip angles responses every 12 hours for 48 hours.

Compared to wild-type, *lazy1* mutant branches grew initially at a wider angle, as reported previously. However, branch tips behaved similarly to wild-type in response to dark conditions, changing their growth trajectory upward, against the gravity vector (Fig. 5B and Supplementary Fig. S2). Under continuous light, *lazy1* mutants had a strong and opposite response, growing downward towards the gravity vector (Fig. 5B and Supplementary Fig. S2), showing a hyper-sensitivity to continuous light. To address whether having a functional *TAC1* gene in *lazy1* mutant plants was necessary for this response to light, we assessed phenotypes of *tac1 lazy1* mutants under these conditions as well. *tac1 lazy1* mutants phenocopied *lazy1* mutants alone, suggesting that *TAC1* is not required for the *lazy1* response to light.

Unlike *lazy1* and *tac1*, single mutants of *dro1, dro2* and *dro3* were reported to have no observable shoot phenotypes, and thus we did not explore the effect of light environments on these single mutants. A *lazy6* single mutant has not yet been described, due to the lack of an available T-DNA insertion line. In a pLAZY6::GUS reporter line, we observed strong *LAZY6* gene expression in the shoot vasculature through 4 weeks of growth (Supplementary Fig. S3A) and engineered a *lazy6* CRISPR mutant to observe its branch angle phenotype. As a single mutant, we saw no substantial difference in shoot architecture from wild-type in light or dark conditions (Supplementary Fig. S3B), and *lazy1 lazy6* mutants showed similar branch angles and branch tip growth orientations (Supplementary Fig. S3B). This suggested that despite the strong differential expression in response to light, loss of *LAZY6* does not significantly impact *lazy1* light response. Thus, we decided to test whether phenotypic changes to branch orientation in response to light required *LAZY/DRO* genes by examining higher order mutants, including the triple mutant containing mutations in *lazy1, dro1*, and *dro3*, and the quadruple mutant containing *lazy1 dro1, dro2*, and *dro3* mutations. Both the triple and quadruple mutants appeared to lose response to continuous dark or light and continued to grow downward with gravity regardless of light condition (Fig. 5D and 5E). We found this to occur in plants whose bolts had grown to at least ∼15cm, while shorter bolts 7-10cm in length continued to grow vertically, or just slightly bent, regardless of light or dark changes. This suggested that multiple IGT genes are required to respond to light cues in order to set branch angles with respect to gravity.

To determine whether light induced changes in *LAZY/DRO* gene expression could conceivably modify light-stimulated changes in branch orientation, we tested if overexpression of *LAZY* and *DRO* genes affected branch growth angle in response to light. To assay this, we overexpressed *LAZY1, DRO1*, or *LAZY6* in the *lazy1* background, and compared the phenotypes in response to light conditions. It was recently reported that expressing *LAZY1* under its native promoter in a *lazy1* mutant background only partially rescued the mutant branch angle phenotype (Yoshihara and Spalding, 2019). We found this to be true when overexpressing *LAZY1* in a *lazy1* mutant background as well (Fig. 6). We observed an upward leaf curling phenotype, an overexpression phenotype described with other IGT genes (Guseman *et al*., 2017; Hollender *et al*., 2020), suggesting that the overexpression construct is at least partially functional. However, constitutive expression of *LAZY1* under the 35S promoter did not fully rescue under continuous light or dark conditions (Fig. 6). Next we overexpressed *DRO1* in a *lazy1* mutant background, and found it not only rescued the branch angle phenotype of *lazy1*, but phenocopied the shoot phenotype conferred by *DRO1* overexpression in both a wild-type and *dro1* mutant background (Fig. 6 and Guseman *et al*., 2017). These lines exhibited narrow, upward growing branches similar to a *tac1* mutant (Dardick *et al*., 2013). Upon growth in continuous light, branches responded by beginning to grow at a wider angle. Growth in the dark, however, had little to no impact, as branch angles were already near vertical (Fig. 6). In contrast to *DRO1, LAZY6* overexpression only partially rescued *lazy1* branch angle phenotype. Branches grown in continuous light responded by growing downward, but not to the extreme of the *lazy1* mutant, and dark grown plants exhibited a phenotype similar to wild-type. These lines did not show a pronounced leaf curling phenotype (Fig. 6).

**Figure 6:**
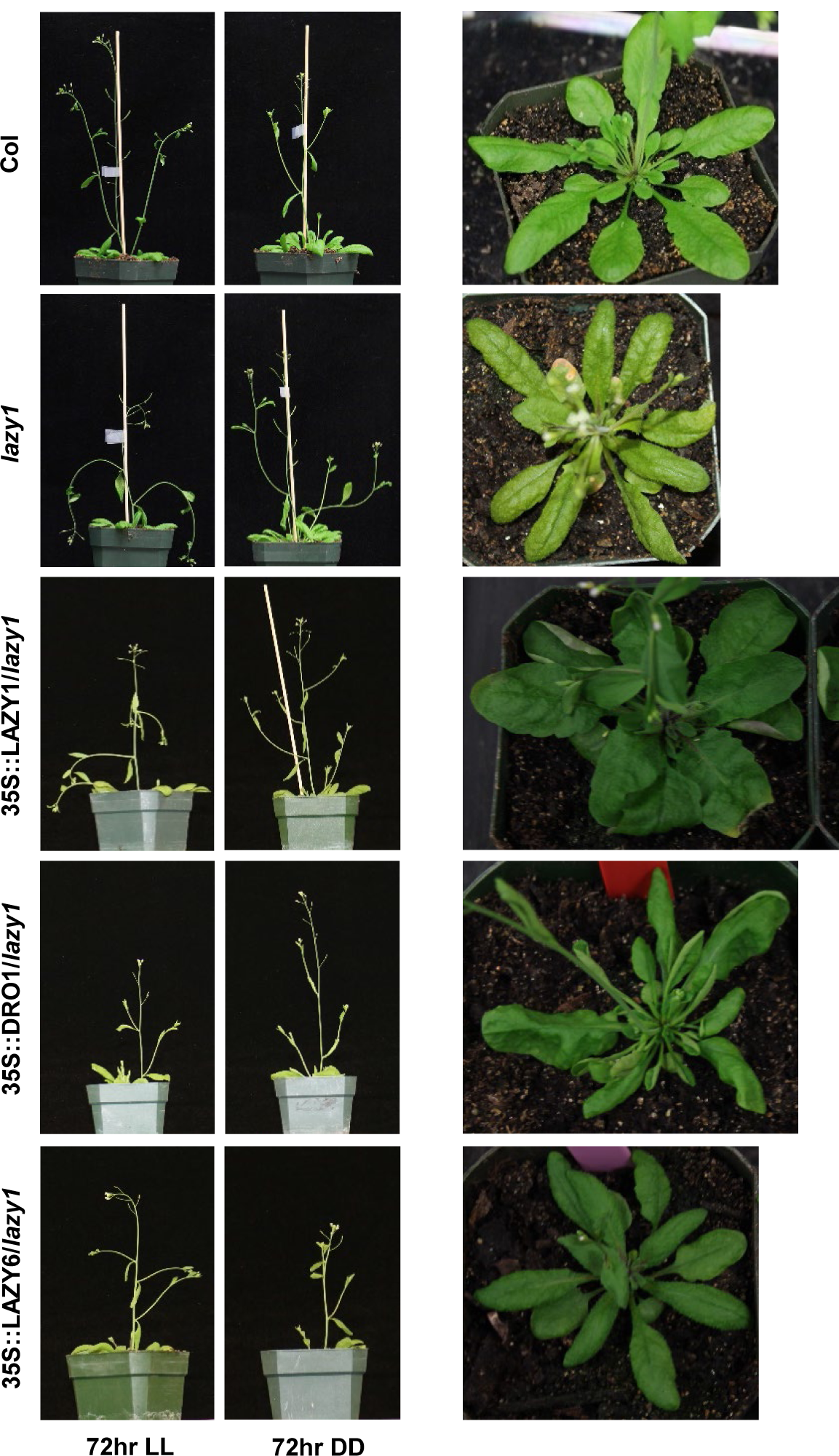
Plants constitutively expressing *LAZY* and *DRO* genes maintain response to changes in light. A. Phenotypes of Columbia wild-type, *lazy1* mutant, and 35S::LAZY1/*lazy1*, 35S::DRO1/*lazy1*, and 35S::LAZY2/*lazy1* overexpression lines after growth in continuous light or dark for 72 hours. B. Images of rosettes representative of these lines demonstrating leaf curling phenotypes in 35S::LAZY1/*lazy1* and 35S::DRO1/*lazy1* overexpression lines.

## Discussion

Manipulation of light, photosynthesis, and circadian inputs revealed a complex picture of environmental regulation of IGT gene expression. By varying light color, light signaling, photosynthesis, and diurnal conditions, we found that different *LAZY* and *DRO* genes have distinct responses to light-related signals. For example, *LAZY1* gene expression was relatively stable throughout changes to light regimes, never changing more than 2-fold in response to varying light stimuli. In contrast, *DRO1* and *LAZY6* showed more dramatic responses to the treatments, particularly circadian experiments, R and B light signaling components, and some photosynthetic treatments, while *DRO2* appeared to have the greatest response to continuous light, both in circadian experiments and change from continuous dark to light. These results suggest that gravity responses mediated by IGT genes are influenced by light-related signals. Thus, these data inform potential experiments for microgravity settings that could tease apart these connections. For example, in microgravity under 16L:8D day length conditions, adult Arabidopsis look strikingly like *lazy1* mutants (Link *et al*., 2014). Phenotyping different *lazy* and *dro* mutants under different light regimes in microgravity may reveal their relationships to light cues and changes.

Loss of photoreceptors and associated signaling pathway components influenced IGT gene expression in a variety of different ways. While *LAZY1* expression showed little to no change in response to growth in R, B, or FR light, its expression increased in both *cry1 cry2* and *phy* mutant backgrounds under multiple light conditions. *LAZY1* expression additionally increased in a *hy5 hfr1 laf1* background and decreased slightly in a *cop1-6* background under some conditions. Together this may suggest *LAZY1* is a downstream target of negative repression by *PHY* and *CRY* signaling. While we have yet to clearly identify a *lazy6* mutant phenotype, *LAZY6* expression showed dynamic and dramatic responses to changes in light and circadian rhythms, and its expression in different photoreceptor-mediated signaling mutant backgrounds often fit with experimentally established R and B light signaling pathways. For example, *COP1* negatively regulates the light signaling integration transcription factors *HY5, HFR1* and *LAF1* downstream of R and B light, and under W light conditions. *LAZY6* was increased in a *cop1-6* mutant background but decreased in the *hy5 hfr1 laf1* mutant. This pattern of *LAZY6* expression changes in R light, where the increase in *cop1-6* is abolished. It is altered in B light as well, where expression increases in both backgrounds. *LAZY6* is strongly upregulated under B light but depends on the presence of both *PHOT* and *CRY* genes. *LAZY6* is also downregulated in R light, and shows a strong increase in phy mutant backgrounds, suggesting negative regulation by this pathway. The *LAZY6* promoter contained several different cis-elements related to phyA signaling and bZip transcription factor binding, which is in line with its strong light-induced changes. *DRO1* and *DRO2* expression responded similarly to R and FR light (both decreased), loss of *cry1 cry2* (decrease), and loss of *hy5 hfr1 laf1* (decrease). While these findings are not necessarily consistent with being targets of the core photoreceptor-mediated signaling pathways, it suggests they have relationships to these signaling pathways and may be similar to each other in the way they are regulated. As *DRO2* is in close proximity with a neighboring gene, it is difficult to determine the length of its promoter and thus its similarity to *DRO1*, however both genes contain several light-related cis-elements, including a phyA-related element in *DRO1*.

Consistent with this, *DRO1* expression also exhibited a strong increase in a *phyAB* and *phyABDE* background under R light, which suggests an additional connection to phytochromes.

It has been reported that *dro1* and *hy5* mutants have very similar root orientation phenotypes, both exhibiting a near-horizontal orientation of root growth (Oyama *et al*., 1997; Guseman *et al*., 2017). Our finding that *DRO1* and *DRO2* expression were significantly and consistently decreased in a *hy5* mutant background (Fig. 5) suggest that *DRO* genes act downstream of *HY5*. Further investigation into their genetic and molecular relationship, as well as DRO1 protein function, may help elucidate these connections.

Photosynthetic inhibitor treatments overall had minor or inconsistent effects, with the notable exception of DCMU treatment in the light, which resulted in significant increases in *DRO1* and *DRO2* expression, about 15-and 2-fold respectively, and about 11-fold downregulation of *LAZY6*. DCMU blocks the electron transport chain via the plastiquinone site of PSII, resulting in ROS production in the light. The changes in *DRO1, DRO2*, and *LAZY6* may suggest a relationship with reactive oxygen pathways. Another notable effect was that of sucrose on *LAZY6* expression. *LAZY6* expression in the dark is dramatically decreased (Fig. 1 and 4), however this was rescued by growth on media containing sucrose, suggesting that dark repression of *LAZY6* may be mediated by photosynthate.

*lazy1* mutants have the strongest single mutant shoot gravitropic set point angle phenotype, but were still able to respond to changes in light. In fact, continuous light caused *lazy1* branch tips to actively grow downward. This appears to be an enhanced response compared to wild-type, which grows at a wider angle under this light condition, but not against the gravity vector. This phenotype is similar to the phenotype observed in triple *lazy1 dro1 dro3* mutants and quadruple *lazy1 dro1 dro2 dro3* mutants under normal (long day) light conditions (Fig. 5 and Yoshihara and Spalding, 2019). While we were unable to obtain or design primers for *DRO3* expression, we did see a dampening of *DRO1* circadian expression peaks (Fig. 2), and total *DRO2* expression under continuous light (Fig. 1 and 2). As these proteins are still intact in a *lazy1* mutant, their dampening and/or loss under continuous light may explain the downward growth phenotype. Alternatively, light may influence the localization or binding of these proteins, affecting the mechanism of their action, however, more work is needed to address this. In continuous dark, *lazy1* branches began to grow upward, and displayed a phenotype similar to wild-type plants, suggesting that *lazy1* has a light-conditional mutant phenotype. Loss of *TAC1* did not appear to alter this re-orientation response, rather *lazy1* appeared to be completely epistatic to *tac1*, which we reported under normal light conditions as well (Hollender *et al*., 2020). However, loss of *DRO1* and *DRO3* in addition to *LAZY1* did lead to a loss of upward re-orientation in continuous darkness (Fig. 5D). After 72 hours in darkness *DRO2* expression increased, while *DRO1* expression decreased (Fig. 1). Altogether, this data is inconsistent with *TAC1* or *DRO1* driving the *lazy1* dark response. While a certain number of IGT genes appear to be required for this dark response, this supports the presence of an additional IGT-independent pathway involved in light responses (Yoshihara and Iino, 2007).

Triple *lazy1 dro1 dro3* mutants exhibit an extreme reverse gravitropic growth defect and show total insensitivity to continuous light and dark regimes, despite containing intact *LAZY5, LAZY6* and *DRO2* genes (Taniguchi *et al*., 2017; Yoshihara and Spalding, 2017). This raises the question of *LAZY* and *DRO* individual genetic contributions to light sensitivity of architectural phenotypes. While a combination of intact *LAZY1, DRO1*, and *DRO3* are required for shoot architecture sensitivity to changes in diurnal regimes, the effect of *DRO2, LAZY5*, and *LAZY6* are less clear. While not exposed to changes in day length, recent experiments show that higher order mutant seedlings containing *dro2* – both *lazy1 dro1 dro2 dro3* and *dro1 dro2 dro3* (*atlazy1,2,3,4* and *atlazy2,3,4* from Yoshihara and Spalding, 2017) have more severe hypocotyl and root gravitropic defect than counterparts lacking the *dro2* mutation, suggesting an essential role in root gravitropism for *DRO2*. In shoots of adult plants, a *lazy1 dro2* mutant appears to have a similar phenotype to *lazy1* alone, and *dro1 dro2* and *dro2 dro3* mutants show little if any shoot phenotype (Yoshihara and Spalding, 2017). However, we were not able to test these mutant’s responses to changes in light, and further study is required to understand how *DRO2* contributes to the light response phenotype. Due to a lack of availability of a *LAZY6* T-DNA insertion mutant, a higher order mutant has only recently been published, and demonstrated that loss of *LAZY6* did not enhance the gravitropic defect of loss of *LAZY1, DRO1, DRO2*, and *DRO3* together in adult shoots (*atlazy1,2,3,4,6* in Yoshihara and Spalding, 2019). Future light experiments with different combinations of higher order mutants will help to clarify the individual roles of these genes in modulating gravity in response to light signals.

While *LAZY* and *DRO* genes are collectively required for response to light, they appear to be functionally diverse. Using constitutive expression lines, we found that *LAZY1* exhibits upward curling leaves, however is unable to rescue the branch angle phenotypes of *lazy1* (Fig. 6). This is consistent with recent work showing that *LAZY1* expression under its native promoter only partially rescues *lazy1* branch angle (Yoshihara and Spalding, 2019). Similarly, *TAC1* expression in a *tac1* mutant background also partially rescued the branch angle phenotype (Dardick *et al*., 2013). In contrast, overexpressing *DRO1* in a *lazy1* mutant shows a leaf curling phenotype, rescues branch angle, and has an upward growth habit similar to *DRO1* overexpression in both *dro1* and wild-type backgrounds. In recent studies, expression of *DRO3* under the *LAZY1* promoter (*pAtLAZY1::AtLAZY2/atlazy1*, Yoshihara and Spalding, 2019) was able to fully rescue the *lazy1* branch angle phenotype, while we found here that *LAZY6* overexpression only partially recues, and does not exhibit leaf curling (Fig. 6). This evidence, along with mutant phenotypes, suggests some level of functional divergence among IGT family proteins, in addition to subfunctionalization in gene expression (Dardick *et al*., 2013; Yoshihara *et al*., 2013; Taniguchi *et al*., 2017; Yoshihara and Spalding, 2017; Guseman *et al*., 2017).

The results presented here add to a growing number of studies revealing how environmental factors influence the ability of plants to sense and respond to changes in gravity and set architecture accordingly (Digby and Firn, 2002; Vandenbrink *et al*., 2014; Roychoudhry *et al*., 2017; Waite and Dardick, 2018). While IGT genes and auxin response have been studied in a variety of different tissues, developmental timepoints, and light conditions, experimental conditions are not standardized, making direct comparisons of results difficult. Our evidence for how light affects these phenotypes demonstrates a need for standardization of environmental conditions in future studies. Further, these studies have largely confounded/equated re-orientation to changes in gravity (fast response) with steady state gravitropic set point angles (slow response). Recent work has demonstrated the dynamics of excised inflorescences from wild-type, *lazy1*, and higher order mutants during the 4-5 hour re-orientation response through the longer >15 hour gravity set point response (Yoshihara and Spalding, 2019). It is worth future study to dissect the roles of these genes in gravity sensing and response.

Together, our findings support a model in which *LAZY* and *DRO* genes are collectively required for branch angle orientation in response to light. The complex expression patterns of IGT family genes suggests that light signals may contribute to branch orientation not only via phototropic pathways but also by modulating gravity responsiveness via differential expression of IGT genes (Fig. 7). Future work will be necessary to uncover the precise roles of individual IGT gene family members in light response, whether this effect is direct or indirect, and whether they are also responsive to other environmental stimuli.

**Figure 7:**
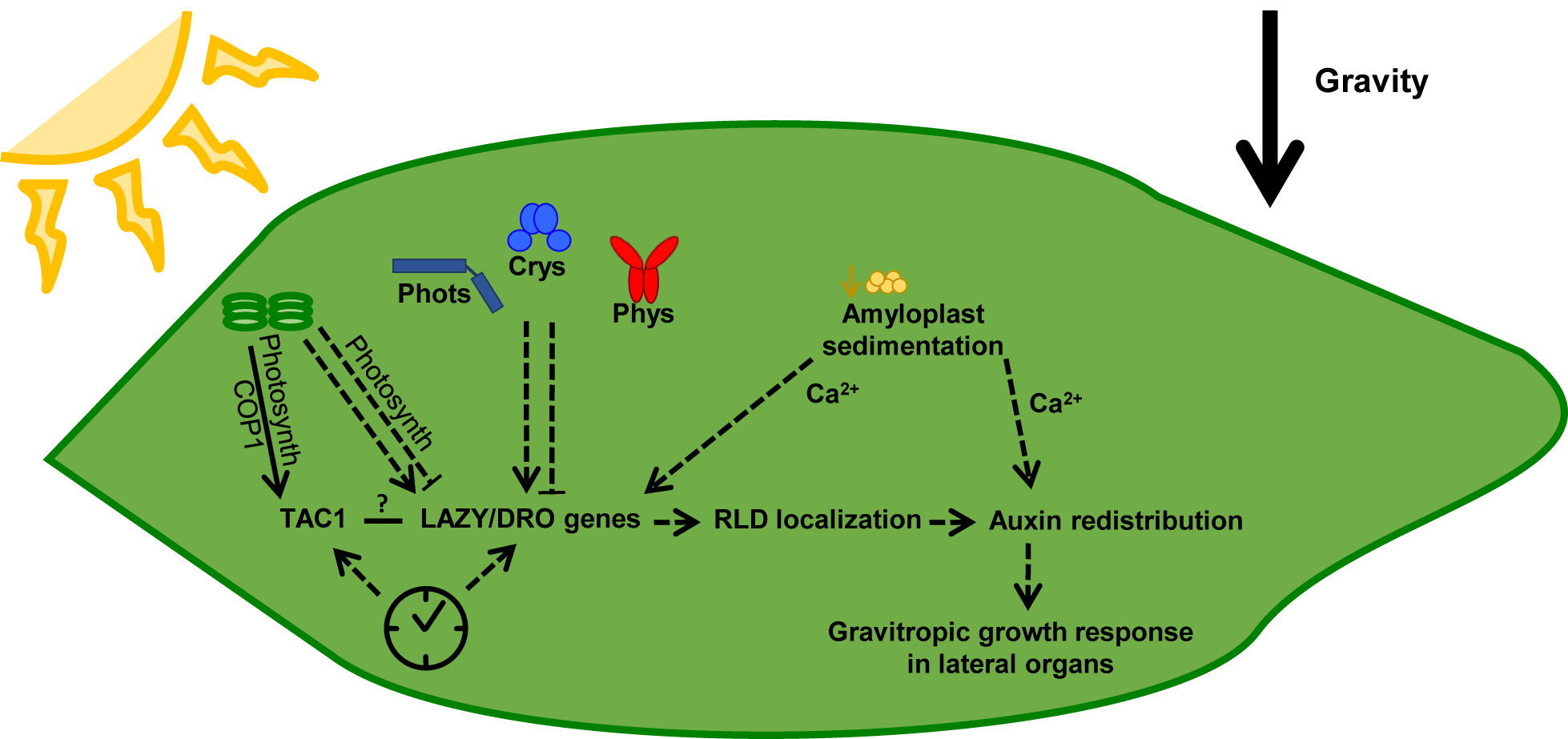
Model for IGT gene integration of light and gravity responses *LAZY* and *DRO* genes are collectively required for branch angle orientation in response to light. *TAC1* is influenced by photosynthetic signals (Waite and Dardick, 2018), while *LAZY* and *DRO* gene expression is differentially influenced by both photoreceptor-mediated signaling components and photosynthetic signals. Most IGT genes additionally show circadian signatures, suggesting influence by the clock. The action of LAZY and DRO proteins is reported to occur downstream of amyloplast sedimentation and upstream of RLD localization and auxin re-distribution in responses to changes in gravity (Taniguchi *et al*., 2017; Furutani *et al*., 2020). Together this supports a model where IGT genes may act as a point of integration between some light signals and gravity response.

## Supplementary Data

Supplementary Table 1: Table of IGT/LAZY family gene names and Arabidopsis IDs from recent studies.

Supplementary Figure 1: *LAZY* and *DRO* genes contain several types of light-related ciselements.

Supplementary Figure 2: Light-driven phenotypic changes of wild-type Columbia and *lazy1* mutants after 48 – 72 hours.

Supplementary Figure 3: *LAZY6* is expressed in shoot vasculature through 28 dpg and does not alter *lazy1* mutant light-response phenotype under constant light or dark.

## Acknowledgements

We would like to thank the labs of Edgar Spalding, Kerry Franklin, Chentao Lin, Ken-ichiro Shimazaki, Xing Wang Deng, Jennifer Nemhauser, and In-Cheol Jang for providing seeds of the light signaling mutants. We thank Courtney Hollender and Edith Pierre-Jerome for critical review of the manuscript. The work at AFRS was supported by Agriculture and Food Research Initiative Competitive grants 2017-67012-26099 and 10891264 from the USDA National Institute of Food and Agriculture and by the National Science Foundation grant number 1339211.

